# Membrane-mediated ligand unbinding of the PK-11195 ligand from TSPO

**DOI:** 10.1101/2020.01.21.914127

**Authors:** Tom Dixon, Arzu Uyar, Shelagh Ferguson-Miller, Alex Dickson

## Abstract

The translocator protein (TSPO), previously known as the peripheral benzodiazepine receptor, is of longstanding medical interest as both a biomarker for neuroinjury and a potential drug target for neuroinflammation and other disorders. Recently it was shown that ligand residence time is a key factor determining steroidogenic efficacy of TSPO-binding compounds. This spurs interest in simulations of (un)binding pathways of TSPO ligands, which could reveal the molecular interactions governing ligand residence time. In this study, we use a weighted ensemble algorithm to determine the unbinding pathway for different poses of PK-11195, a TSPO ligand used in neuroimaging. In contrast with previous studies, our results show that PK-11195 does not dissociate directly into the solvent but instead dissociates via the lipid membrane by going between the transmembrane helices. We analyze this path ensemble in detail, constructing descriptors that can facilitate a general understanding of membrane-mediated ligand binding. We construct a Markov state model using additional straightforward simulations to determine pose stability and kinetics of ligand unbinding. Together we combine over 40 *µ*s of trajectory data to form a coherent picture of the ligand binding landscape. We find that all poses are able to interconvert before unbinding, leading to single mean first passage time estimate for all starting poses which roughly agrees with the experimental quantity. The ligand binding transition state predicted by our combined model occurs when PK-11195 is already in the membrane and does not involve direct ligand-protein interactions. This has implications for the design of new long residence-time TSPO ligands.

**SIGNIFICANCE:** Kinetics-oriented drug design is an emerging objective in drug discovery. However, while ligand binding affinity (or the binding free energy) is purely a function of the bound and unbound states, the binding kinetics depends on the nature of the paths by which the (un)binding occurs. This underscores the importance of approaches that can reveal information about the ensemble of (un)binding paths. Here we used advanced molecular dynamics approaches to study the unbinding of PK-11195 from TSPO and find it dissociates from the protein by dissolving into the membrane, and that the transition state occurs after the PK-11195 molecule has already separated from TSPO. These results motivate the design of future long-residence time TSPO ligands that destabilize the membrane-solvated transition state.

## INTRODUCTION

The binding affinity of a ligand to its protein target has long been viewed as the key parameter determining its efficacy. However, recent studies have shown that in some protein-ligand systems residence time (RT) correlates more strongly with efficacy than binding affinity (1). But unlike the binding affinity, RT is not a state function; it depends on the height of the free energy barrier separating the bound and unbound states. In order to rationally design ligands for longer RTs we need to understand the (un)binding mechanism and what molecular interactions occur along the ligand (un)binding pathway.

Previous studies have shown that the translocator protein 18kDa (TSPO) is one such protein where RT is important for predicting efficacy (2). TSPO is a well-conserved membrane protein, being present all kingdoms including prokaryotes as well as in the outer mitochondrial membrane of eukaryotes (3). TSPO has five transmembrane *α*-helices (TM1-5) along with a small helical region in a 20-residue loop connecting TM-1 and TM-2 on the cytosolic side (Fig. 1A). While in the membrane, TSPO is largely found in a dimeric state (4). To date, four different structures have been solved for TSPO, for both bacterial (4, 5) and mammalian (6, 7) organisms the former by X-Ray crystallography, the latter by NMR.

**Figure 1:**
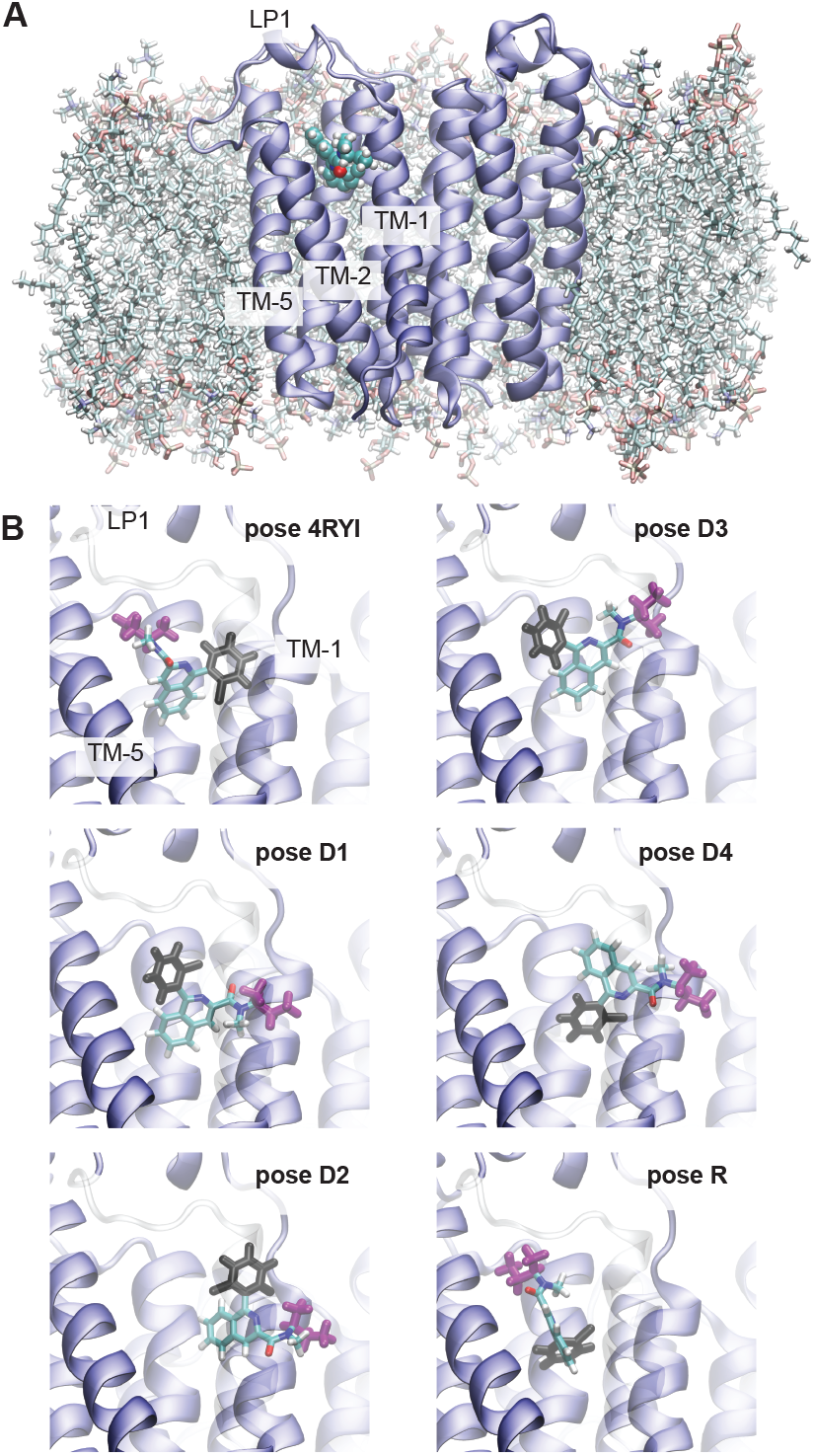
TSPO-PK11195 system. (A) Front view of the TSPO dimer in the membrane with PK bound. (B) All six starting poses are shown from the side view, along the inter-dimer axis. To compare poses, two moeities of PK are colored in black (o-chlorophenyl) and magenta (1-methylpropyl), with the rest of the molecule colored according to atom name. TM-2 is shown as transparent for clarity.

While the structure of TSPO have been solved, its function remains unknown. In humans, TSPO is highly expressed in steroidogenic tissues, leading to the hypothesis that it is involved in the regulation of cholesterol transport across the mitochondrial membrane.

Indeed, TSPO has been shown to have a high binding affinity for cholesterol (8). There are other studies linking it to apoptosis (9, 10) and cellular stress regulation in TSPO knockout mice (11, 12), although evidence for this is mixed (13, 14). Increased TSPO expression has also been observed in cases of neurodegenerative diseases such as Alzheimer’s and Parkinson’s diseases (15). Relatedly, due to its high expression in areas of inflammation TSPO serves as a biomarker for neurodegenerative disease and brain trauma, and radiolabeled ligands such as [H3]-PK-11195, are commonly used in positron emission tomography (PET) scans (16). PK-11195 (hereafter denoted “PK”) is an isoquinoline carboxamide with no known therapeutic effect (14) and a RT of 34 min (2, 17).

Molecular dynamics (MD) simulations have been previously performed using a bound TSPO-PK complex. Researchers recently determined the unbinding pathway of PK from a rat TSPO model generated from the PDB 2MGY structure (18). To generate unbinding paths they used a combination of random accelerated MD (RAMD) (19) and steered MD (20) and determined that PK unbinds into the cytosol through the largely disordered LP1 region (Fig. 1A). Unfortunately, this starting structure, determined by NMR, was significantly destabilized by the detergent used in the purification (21, 22). Also, the methods used to determine the unbinding pathway RAMD have the potential to impart bias on the predicted (un)binding path. Another group performed an induced-fit docking of PK using Glide (23) with a homology model to resemble the mammalian (mouse) TSPO structure using the PDB 4UC1 *Rhodobacter sphaeroides* structure. They simulated the TSPO-PK complex for 700 ns and did not observe significant ligand displacement, which is expected due to the extremely long RT of the TSPO-PK complex.

Here we study the unbinding mechanism for the TSPO-PK complex, using PDB 4UC1 as the TSPO starting structure (4) and using a weighted ensemble algorithm: Resampling of Ensembles by Variation Optimization (REVO) to generate continuous unbinding pathways without perturbing the underlying dynamics (24). REVO has been previously applied to study ligand unbinding on a series of host-guest systems (25) and the trypsin-benzamidine system (24). We identify key residues which interact strongly with PK as it dissociates from TSPO. We finally compute the residence times for PK unbinding at each pose and compare these results to experimentally determined residence times.

In the next section we discuss the methodology used for the simulations, the REVO resampling algorithm, the clustering algorithm used to make the CSN, and rate calculations. The results are presented in Results and Discussion, including the pathways found for dissociation of PK from TSPO, residues which bound strongly to PK along the observed pathways, and comparison of residence times between different poses. We then summarize our findings and discuss how they relate to the existing research.

## MATERIALS AND METHODS

### Protein Preparation

The initial translocator protein (TSPO) dimer structure is comprised of chains A and B from PDB 4UC1(4). This x-ray crystal structure comes from the *Rhodobacter sphaeroides* with an A139T mutation to resemble human TSPO. CHARMM-GUI membrane generator(26) was used to place the TSPO complex into a membrane comprised of 174 phospholipids consisting of 53.4% phosphatidylcholine, 28.2% phosphatidylethanolamine, and 18.4% phosphatidylinositol lipids. 10268 TIP3 water molecules were inserted up to a cutoff of 10 Å from the complex and 121 potassium ions and 27 chloride ions were added to reach a salt concentration of 150 mM and to neutralize the system. The system was placed into a rectangular box with dimensions 96.4 Å x 96.4 Å x 91.8 Å. The protein was simulated using the CHARMM36 forcefield (27) and parameters for the PK-11195 ligand were obtained with CGenFF (28, 29).

### Docking

Six different PK-11195 (PK) poses were used in the simulations. Docking was carried out with High Throughput Virtual Screening (HTVS) by using Schrödinger Glide(30). The center of mass (COM) of PK-11195 (PK) was placed at the COM of the bound Protoporphyrin IX in the chain A monomer of TSPO protein from PDB 4UC1 without any constraints. PK poses above a threshold Glide GScore (6.5 kcal/mol) were analyzed in post-docking applications of Maestro(31). A homology model of PK-bound TSPO (Pose R) was generated by Xia *et al*. as a Rosetta comparative model of the mouse TSPO structure constructed using TSPO structures from *Mus musculus* (PDB 2MGY(6)), *R. sphaeroides* (PDB 4UC1(4)), and *Bacillus cereus* (PDB 4RYI (5)); more details found in Ref. (32). The TSPO monomer bound to PK from this model was then aligned to chain A of the 4UC1 structure using PyMol 1.7.2.1 (33), and the ligand coordinates from the D1 pose were changed to reflect the new pose. The 4RYI pose was generated by X-Ray crystallography and the coordinates of the PK ligand were added to the 4UC1 structure in the same way as pose R. The system’s energy was minimized using a series of constraints with scripts provided by CHARMM-GUI for all poses. The molecular structure for each pose is shown in Fig. 1B

### Molecular Dynamics

All molecular dynamics (MD) simulations were performed using OpenMM(34) v7.1.1. The timestep for every simulation was 2 fs. To enforce constant temperature and pressure, a Langevin heat bath was used with a set temperature of 300K and a friction coefficient of 1 ps^-1^ was coupled to a Monte Carlo barostat set to 1 atm and volume moves were attempted every 50 timesteps. The non-bonded forces were computed using the CutoffPeriodic function in OpenMM with a cutoff of 10 Å. The atomic positions and velocities are saved every 15, 000 timesteps, or every 30 ps of simulation time, which is the resampling period (*τ*) used here.

### REVO Resampling

To observe long timescale unbinding of PK, we used a variant of the weighted ensemble algorithm: Resampling of Ensembles by Variation Optimization (REVO)(24). In this algorithm, we perform unbiased MD simulation on 48 separate trajectories in a parallel fashion. Each of these trajectories (called “walkers”) has a statistical weight (*w*) that governs the probability with which it contributes to statistical observables. With periodicity *τ*, a resampling procedure is performed, where similar walkers are merged together and unique walkers are cloned, as defined by a distance metric. During cloning, weights are split, and during merging, weights are added, to ensure conservation of probability.

Below we briefly describe the REVO method, focusing on the details of its application in this work. More information on the algorithm can be found in previous work (24). In REVO, merging and cloning is done to maximize a variation function: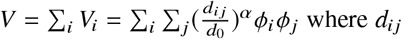 where *d*_*ij*_ is the distance between walker i and walker j determined using a distance metric of choice. For these simulations the distance metric used was the root mean square deviation (RMSD) of the PK atoms between each walker, following alignment to a selection of binding site atoms in TSPO. The exponent *α* is used to modulate the influence of the distances in the variation calculation and was set to 4 for all simulations. *d*_0_ = 0.148 nm is a characteristic distance used to make *V* dimensionless and to normalize the variance for comparison between different distance metrics. *ϕ* is a novelty and here is defined as: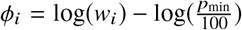. The minimum weight, *p*_min_, allowed during the simulation was 10^−12^. The walker that is selected for cloning is the one that has the highest *V*_*i*_ and the resultant weight of the clones is larger than *p*_min_. The two walkers selected for merging are at most 2 Å away, have a combined weight lower than the maximum allowed weight *p*_max_ = 0.1, and is the walker pair *j, k* that minimizes 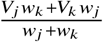. Once the walkers (*i, j, k*) are selected, the new variation is calculated: if it increases, then these operations are performed and another *i, j, k* is proposed; if it decreases then resampling for that cycle is terminated and a new cycle of MD is performed. Three simulations were run for each docked pose using 48 walkers and 1200 cycles, for 1.728 *µ*s of simulation time per simulation. In total each pose was simulated for 5.184 *µ*s.

### Boundary Conditions

The overall goal of the simulations was to determine the pathways along which PK can transition from the initial starting poses to an unbound state. During the simulations, we defined PK as being unbound when the minimum distance between the ligand and TSPO was at least 10 Å. When the ligand crossed this boundary, the weight is recorded and the walker was “warped” back to the initial conformation. The structure recorded before warping is known as an exit point. When the walker warps back, the atomic positions and velocities are reset to their initial values before the simulation began. The walker weight does not change as a result of warping.

### Clustering and Network Layout

The trajectory frames of all 18 REVO runs were clustered together using the MSMBuilder 3.8.0 python library. The frames were featurized using a vector of atomic distances between TSPO and PK atoms initially within 8 Å of each other from the 4RYI starting pose for a total of 7527 distances. A k-centers clustering algorithm was used to generate 2000 clusters using the featurized space and each frame was assigned to a cluster. The clustering was done using the Canberra distance metric. A count matrix describing the cluster-cluster transitions was calculated for a lag time of 30 ps.

We then construct a Conformational Space Network (CSN) from the count matrix, which is a graphical representation of the transition matrix. Each node, representing each row of the transition, and the edges, representing non-zero off diagonal elements of the transition matrix, were determined using the CSNAnalysis package (35). Gephi 0.9.2(36) was used to visualize the CSN. The size of each node is proportional to the statistical population of the cluster. For visualization, the smallest node was set to be 20 times smaller than the largest node. The layout of the network was determined using a force minimization algorithm, Force Atlas included in Gephi. The algorithm repulses nodes that are not connected and attracts nodes that are connected via an edge. The strength of the attractive force is proportional to the weight of the edges. The directed edge weights were values between 0.1 and 100 as determined by *w*_*ij*_ = 100*p*_*ij*_, where *p*_*ij*_ is the transition probability of cluster *i* transitioning to cluster *j*. Unidirectional edge weights were then determined using the average between the two directed edge weights. Force Atlas was applied twice. The first minimization was done without adjusting for node sizes, allowing the nodes to overlap. The second minimization adjusted for the node size and prevented overlap. For visualization, all edges are shown with a uniform line weight.

### Quantifying Unbinding Pathways

Upon analysis of the simulation results, the only unbinding pathways observed in our simulations were PK dissociating through pairs of transmembrane helices. We therefore introduce the coordinate *Q*_*ij*_ which measures the minimum *x*-*y* distance from the COM of PK to the line formed by the COMs of helices *i* and *j* to measure the dissociation progress of PK into the membrane. Negative values indicate the COM of the ligand is closer to the center of the helical bundle, and positive values indicate the COM is closer to being fully dispersed in the membrane. All six poses had trajectories where PK traveled between transmembrane helices 1 and 2 and only pose R had trajectories where PK went between transmembrane helices 2 and 5. For pose R analysis we separate the conformations according to which value (*Q*_12_ or *Q*_25_) is largest. Projections onto a given *Q* value will only use conformations for which that *Q* value is the largest.

### Calculating Non-bonded Energies

We calculated the non-bonded interaction energies by *E*_*int*_ = *V*_*LJ*_ + *V*_*ES*_ where *V*_*LJ*_ is the Lennard-Jones potential energy and *V*_*ES*_ is the potential energy from electrostatic interactions. The Lennard-Jones interactions were determined using a 12 – 6 potential given as: 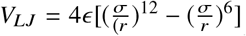 where *r* is the atomic distance between atoms, *σ* is the inter-atomic distance at which the potential is 0, and *∊* is the depth of the potential well. To calculate *σ* and *∊* we used the Lorentz-Berthelot combining rule. There was a hard cutoff distance of 10 Å when calculating the Lennard-Jones potential. The electrostatic energy was calculated using: 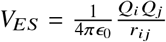 where *Q*_*a*_ is the charge of atom *a, r*_*ij*_ is the interatomic distance between atoms *i* and *j*,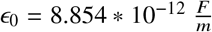 is the permittivity of free space in farads per meter. The specific *σ, ∊*, and *Q*, for each atom type was provided by CHARMM-36 parameter files obtained through CHARMM-GUI. Two sets of non-bonded energies were calculated: between PK and TSPO, and between PK and lipids in the membrane.

### Calculating Off-Rates and Mean First Passage Times using Hills Relation

The rates are calculated using the flux of trajectories into the unbound basin, also known as the Hill relation(37–39), defined as 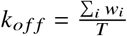 where *w*_*i*_ is the weight of the walker entering the unbound basin, and T is the total simulation time. During the simulations the unbound basin was defined by the 10 Å boundary condition. However, although many walkers had dissociated into the membrane, no walkers made it to the boundary. Therefore, to obtain estimates of unbinding rates, after the simulations were completed the unbound basin was redefined using a minimum distance of 5 Å. In our simulations we observed a total of 2285 instances of trajectory crossings into the 5 unbound basin. This is broken down by starting pose as follows: 4RYI (47), D1 (4), D2 (1804), D3 (278), D4 (152) and R (0). In our analysis, once a walker entered the unbound basin, we ignored all future trajectories associated with that walker. This was done to prevent double-counting of unbinding transitions. The mean first passage time (MFPT), synonymous with the residence time, was calculated as 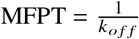. The uncertainty of off rates and MFPT for each pose is the standard error across each set of simulations.

### Calculating Mean First Passage Times using Markov State Models

We create transition matrices, *T* (*τ*), for various lag times (*τ*) using the cluster identities from the CSN and tracking walkers through merging and cloning operations in the REVO resampler. We alter these matrices to include a probability sink for states that are unbound, defined as when PK is at least 5 Å away from the TSPO dimer. We run a Markov chain simulation for a given starting pose and lag time by initializing a probability vector, *P*, where all of the probability starts at the state of a given starting pose. To progress the simulation we use the following: *P*_*k*_ = *P*_0_*T τ* ^*k*^ where *P*_0_ is the initial probability vector, and *P*_*k*_ is the probability vector after *k* timesteps. We continue the simulations until all the probability accumulates in the unbound basin. We then calculate the MFPT using the following formula: 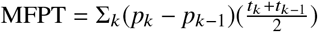 where *p*_*k*_ is the probability of being unbound at timestep *k* and *t*_*k*_ is the time associated with timestep *k*. We repeat this for all initial poses and lag times to determine MFPT as a function of lag time.

### Selecting Poses for Straightforward MD Simulations

To strengthen the accuracy of our Markov state model, we run straightforward simulations at weak points in the network. To determine these weak points, we randomly multiplied the elements of a row on the transition matrix with numbers drawn from a Gaussian distribution with a mean (*µ*) at 1 with a standard deviation (*σ*) of 0.2 and we renormalized the row after perturbation. We rerun the Markov chain simulations to calculate the MFPT. To get a sense of how consistently the cluster alters the MFPT, we randomly perturb the transition matrix 10 times independently. Weak points in the network are determined by the clusters whose perturbations affect the MFPT the most, using the following formula: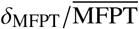, where *δ*_MFPT_ and 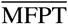 are the standard deviation and average of the perturbed MFPT values, respectively. For two poses, this ratio was greater than 0.2; we identified these clusters as weak points and reran straightforward MD simulations from the highest weighted structure in that cluster. From each weak point we launched 144 independent straightforward MD simulations for a length of 500 cycles (15 ns). In addition we launched trajectories from high-LASA (lipid accessible surface area) clusters in the central unbound region and each of the high-LASA states originating from pose R. In total we ran 10.8 *µ*s of supplemental trajectories to bolster our Markov state model.

## RESULTS

### PK11195 Unbinding Pathway

We comprehensively studied the TSPO-PK interaction landscape using a set of REVO simulations initialized at six different starting poses (Fig. 1B), simulating 5.184 *µ*s per pose. After the simulations were completed, all frames were clustered together into a CSN shown in Fig. 2, where each node represents a PK pose and the edges reveal which poses interconvert in our simulations within a 30 ps lag time. All of the starting poses form a connected network, though pose R is only connected via two low probability edges to the pose 4RYI ensemble (Fig. S2). The 4RYI pose is similarly connected to pose D4, but is also connected to the other docked poses via the high lipid accessible surface area (LASA) clusters. It is worth noting that both pose 4RYI and pose R were the only poses that were not designed for this specific protein structure and were instead inserted from other protein structures after alignment. Consistent with this fact, both of these regions in the CSN do not show accumulation of probability into one or more high-probability states. Instead we observe a broader distribution among many low probability states indicating a lack of a local funneling in the energy landscape. Interestingly, all of the docked poses (D1-D4) show at least one high-probability state, although this is not necessarily at the initial docked pose itself, indicating that some relaxation is required from the docked poses to reach the true local minima.

**Figure 2:**
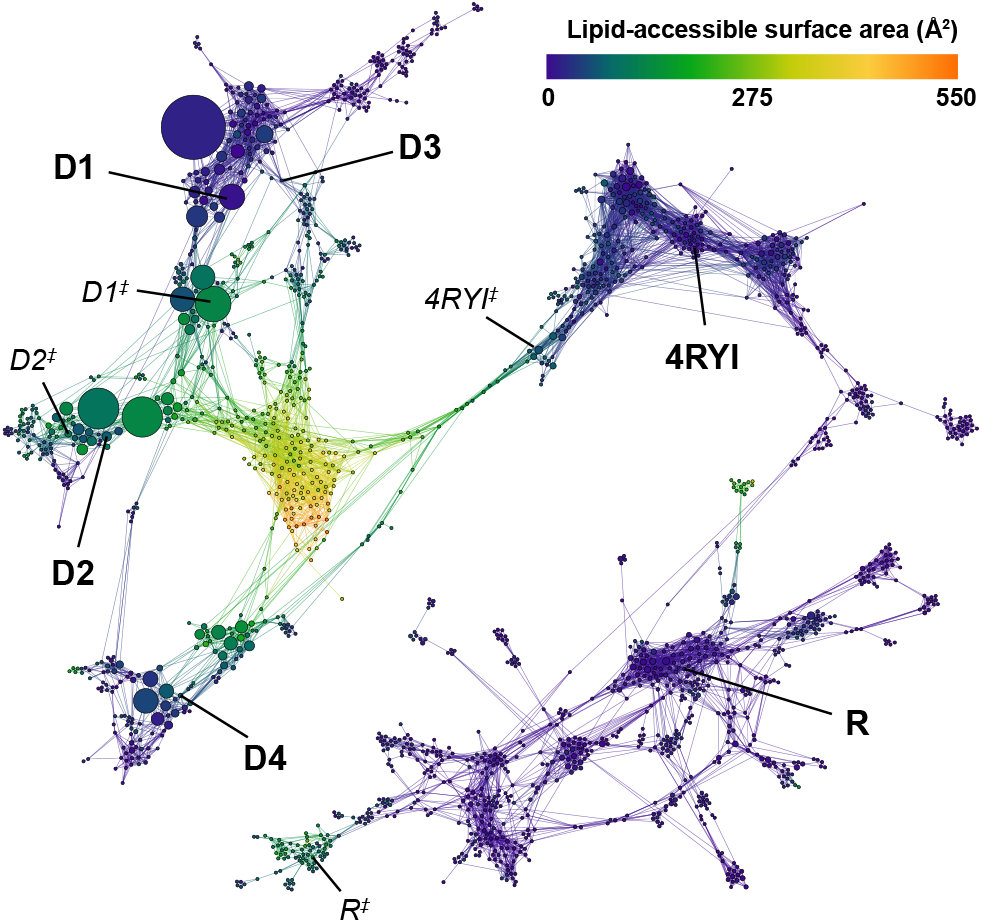
Combined CSN of all REVO simulations from each starting pose. Each node in the network represents a cluster of ligand poses and is sized according to the cluster weight. Nodes are connected by edges if the ligand poses are observed to interconvert in the REVO trajectory segments. Nodes are colored according to the lipid-accessible surface area (LASA). Starting poses are marked in bold and transition state poses shown in Fig. 3D are marked in italics.

Contrary to what was observed in previous work (18), PK did not dissociate into the solvent via the LP1 region: it instead dissociates into the membrane. The CSN shows that all the poses, besides pose R, connect directly to the unbound states, shown in yellow and orange, where PK is fully dissolved into the lipid membrane. In all of these pathways, PK exits between TM-1 and TM-2. The pose R trajectories show two different pathways that have a moderate LASA – one between TM-1 and TM-2 and another between TM-2 and TM-5 – where PK forms direct interactions with membrane lipids.

We introduce the coordinate *Q*_*ij*_, which measures the minimum *x*-*y* distance from the center of mass of the ligand to the line connecting the centers of mass of helix *i* and helix *j* (Fig. 3A). Negative *Q* values indicate the ligand is within the helical bundle and positive values indicate the ligand is outside the bundle. This provides a basis to compare between different pathways and a means of obtaining general information about membrane-mediated ligand unbinding pathways. Fig. 3B compares the TSPO-PK interaction energy (*E*_*int*_) with membrane-PK interaction energy. In the *Q*_12_ pathway (solid lines), PK interacts more closely with the lipid membrane than TSPO after about 5 Å. For the *Q*_25_ pathway (dashed lines) this crossover occurs at 7.5 Å. The difference is due to differences in the orientation of PK along the two pathways. Fig. 3D shows the transition states labelled in Fig. 2 where the *Q* values are approximately equal to zero along each dissociation pathway. We see that each structure is still heavily informed by its starting pose, with very different PK orientations. Fig. 3C shows probability distributions projected onto *Q*_12_ for starting poses D1 and D2. This shows that although D1 started further backward on the unbinding pathway, the simulations discovered another high-probability basin around *Q*_12_ = 0, which can also be seen by the high-probability states around D1^‡^. A representative *Q*_12_ dissociation pathway is shown and analyzed in Fig. 3E and 3F. Note that while the *Q*_12_ value increases steadily along the pathway, the minimum distance between TSPO and PK (used to define the unbound state) rises rapidly only as PK reaches a *Q*_12_ of about 15 Å.

**Figure 3:**
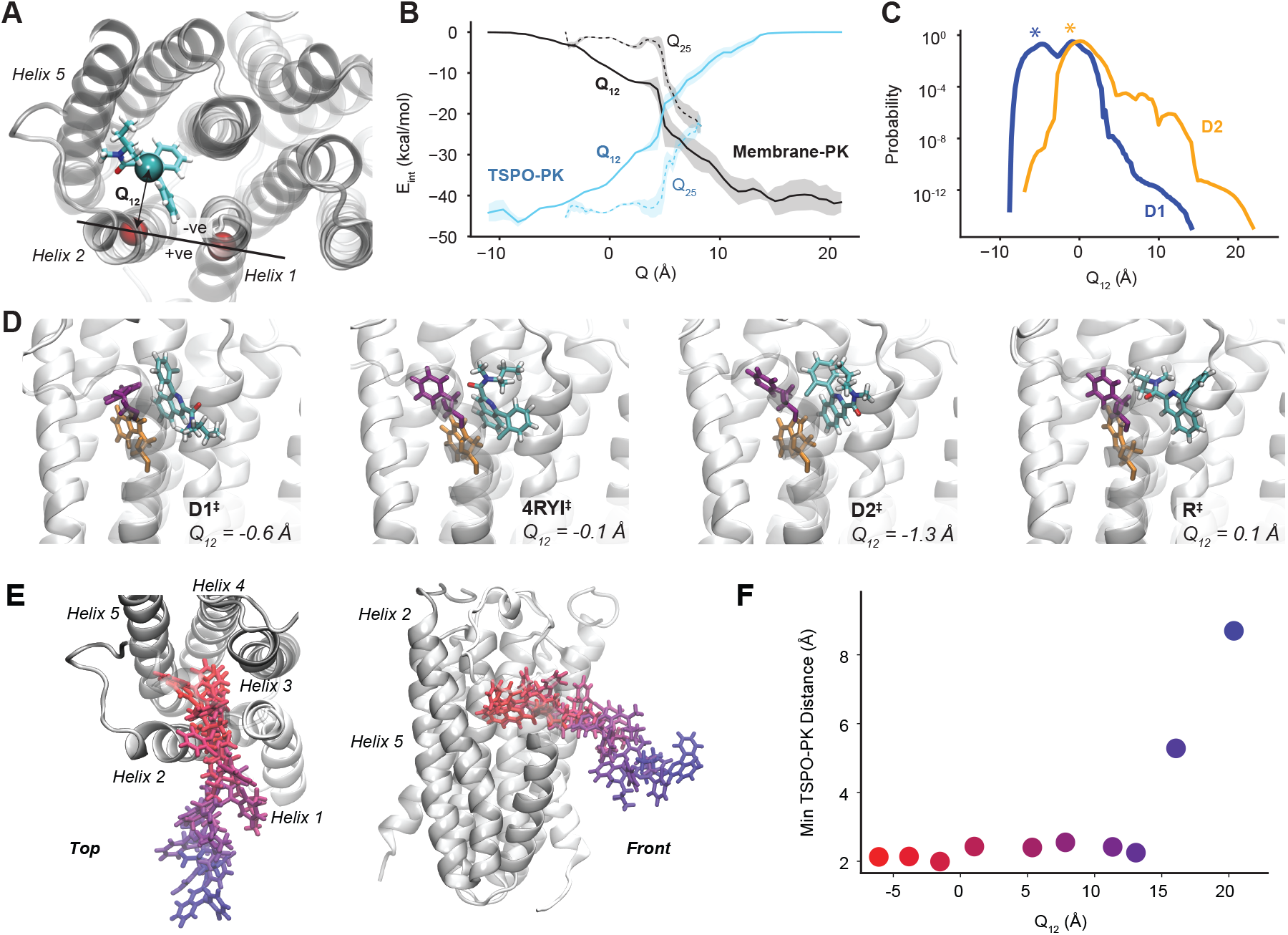
Analysis of membrane-mediated exit paths. (A) The coordinate *Q*_*ij*_ is defined as the *x*-*y* distance between the center of mass of PK, shown as sticks and colored by atom type, and the line that connects the centers of mass of helix *i* and helix *j*. LP1 is not shown here for clarity. (B) The expectation values of the interaction energy between PK and TSPO (blue) and between PK and the membrane (black) are shown as a function of *Q*. In each case the solid line shows *Q*_12_ and the dashed line shows *Q*_25_. The shaded region indicates the standard error over the ensemble of measurements at each *Q* value. (C) Probability curves projected onto *Q*_12_ for simulations initialized in Pose D1 (blue) and D2 (orange). *Q*_12_ values of the starting structures are marked with (*). (D) Poses from transition pathways with *Q* ≈ 0. These poses are also labeled in the CSN of Fig. 2. Phe46 is shown in purple and Trp50 is shown in orange. (E) A set of poses along the *Q*_12_ pathway colored from bound (red) to unbound (blue). Top view is shown on the left and a front view is shown on the right. (F) The minimum PK-TSPO distance and the *Q*_12_ value is shown for each pose in panel (E).

We also measure interaction energies between PK and individual residues for all residues on TM1, TM2, TM5 and LP1 (Fig. S4-S7). Early in both the *Q*_12_ and *Q*_25_ pathways, PK strongly interacts with aromatic residues Phe46 and Trp50 forming *π*-*π* interactions. These aromatic residues with long sidechains follow PK along the unbinding pathway, which is observed by plotting the *Q* value of individual residues as a function of *Q*-PK (Fig. S8 and S9). Interestingly, this phenomenon occurs for smaller amino acid sidechains as well: Gly22 and Pro47 both change *Q* value significantly over the *Q*_12_ pathway, indicating significant distention of the helices during unbinding.

### PK11195 Rates and Residence Times

We directly estimate the unbinding rates (*k*_*of f*_) by summing the weights of the unbinding trajectories and we calculate the mean first passage time (MFPT) by inverting the unbinding rate for each starting pose (Fig. 4A). Pose D2 had a high unbinding flux and a predicted MFPT of less than 0.02 s, indicating a clear lack of stability with respect to the other poses. Poses D3 and D4 had predicted MFPTs of 2.6 and 4.1 minutes, respectively, still lower than the experimental measurements; these estimates are likely to continue to decrease with further simulation time. Poses 4RYI and D1 had MFPT estimates near or above the experimental MFPT (28 and 260 min, respectively). No unbinding events were observed for Pose R, implying an even longer MFPT than 260 min.

**Figure 4:**
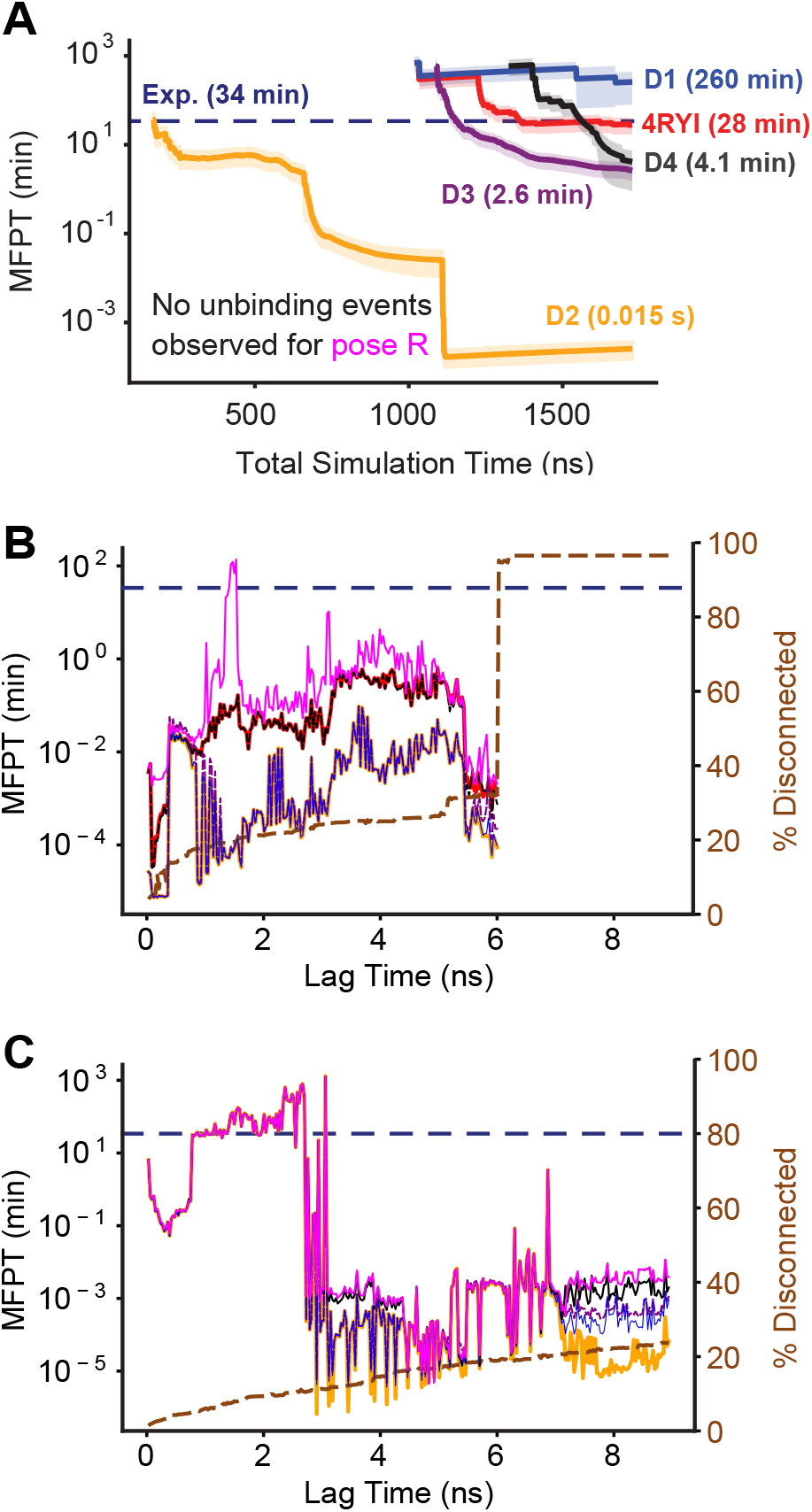
Convergence of mean first passage time (MFPT) estimates using (A) unbinding fluxes using REVO simulations and the Hill relation, (B) a Markov state model constructed with trajectory segments from the REVO simulations and (C) a Markov state model constructed with trajectory segments from the REVO and straightforward simulations. The light shaded area in (A) shows the standard error across the three simulations conducted for each pose. Poses 4RYI and D4, and poses D1, D2, and D3 had nearly identical MFPTs. The brown dashed line shows the percentage of states that were disconnect from the main network The experimental MFPT of 34 min is shown as a dashed blue line in each panel.

One of the issues with performing simulations via weighted ensemble is ensuring the simulations converge. The lack of convergence introduces additional uncertainty into *k*_*of f*_ and MFPT calculations. To address this issue, we launch a set of Markov chain simulations using the transition matrix that constructed the CSN as well as transition matrices at longer lag times (see Section). The results show that all poses except pose R are several orders of magnitude below the experimentally determined residence time (2) of 34 ± 3 for all lag times. For most lag times, the pose R MSMs also predict a lower MFPT by several orders of magnitude, but at a lag time of 1.75 ns, the MFPT briefly rises into the minute range (Fig.4 B). Additionally, the D1, D2, and D3 MFPTs are almost identical, suggesting that these poses interconvert in the MSM. Similar results are seen between poses 4RYI and D4. By tracking the probability of being in a specific pose over the course of a Markov chain simulation, we see this is indeed the case (Fig. S12). For pathways starting in either D1, D2, or D3, we observe that before they reach the unbound basin, the probability distributions form the same quasi-equilibrium mixtures between the D1, D2 and D3 ensembles, where D3 is the most probable and D2 is the least probable. Unbinding pathways starting in D4 and 4RYI do not visit the D1-D2-D3 states. They instead form their own quasi-equilibrium along the unbinding pathway, with D4 being more probable than 4RYI. Simulations starting from pose R also join the D4-4RYI quasi-equilibrium, but have a longer MFPT as pose R is only connected to the rest of the network through 4RYI.

The accuracy of the MFPT calculations however, assumes that the transition matrix determined from the simulations has converged. However, at different lag times the MFPT for each starting pose can vary wildly, indicating that the transition matrix is sensitive to the inclusion or exclusion of individual trajectory segments. To identify these sensitive transition matrix elements, we perturb individual rows of the transition matrix and re-ran the Markov chain simulations for each starting pose. We repeated this 10 times to test the sensitivity of each cluster to the MFPT calculations. We found three clusters that significantly altered the MFPT, two that were directly connected to one another and attached pose R to the rest of the network, and a third that connected pose D4 to pose D2. Intuitively, in each of these cases the clusters were in regions of the CSN that were sparsely populated, and acted as bottlenecks that connect large regions of the network together.

After identifying the states in the CSN that significantly affect the estimated MFPT, we ran straightforward simulations from the highest weighted frame from the REVO simulations that was clustered in that state. We also ran straightforward MD simulations from high-LASA poses in the *Q*_12_ and *Q*_25_ pathways that were seen by pose R as well as the most probable state in the unbound basin. We then reclustered and remade the CSN network to include the new frames (Fig. 5). Several connections were formed between pose 4RYI and pose R, which also gained connections the other poses after reclustering. Additionally, the most probable region in the network was once again the D1-D3 basin as determined by the steady state probabilities of each state, indicating that D1 and D3 are the most stable poses.

**Figure 5:**
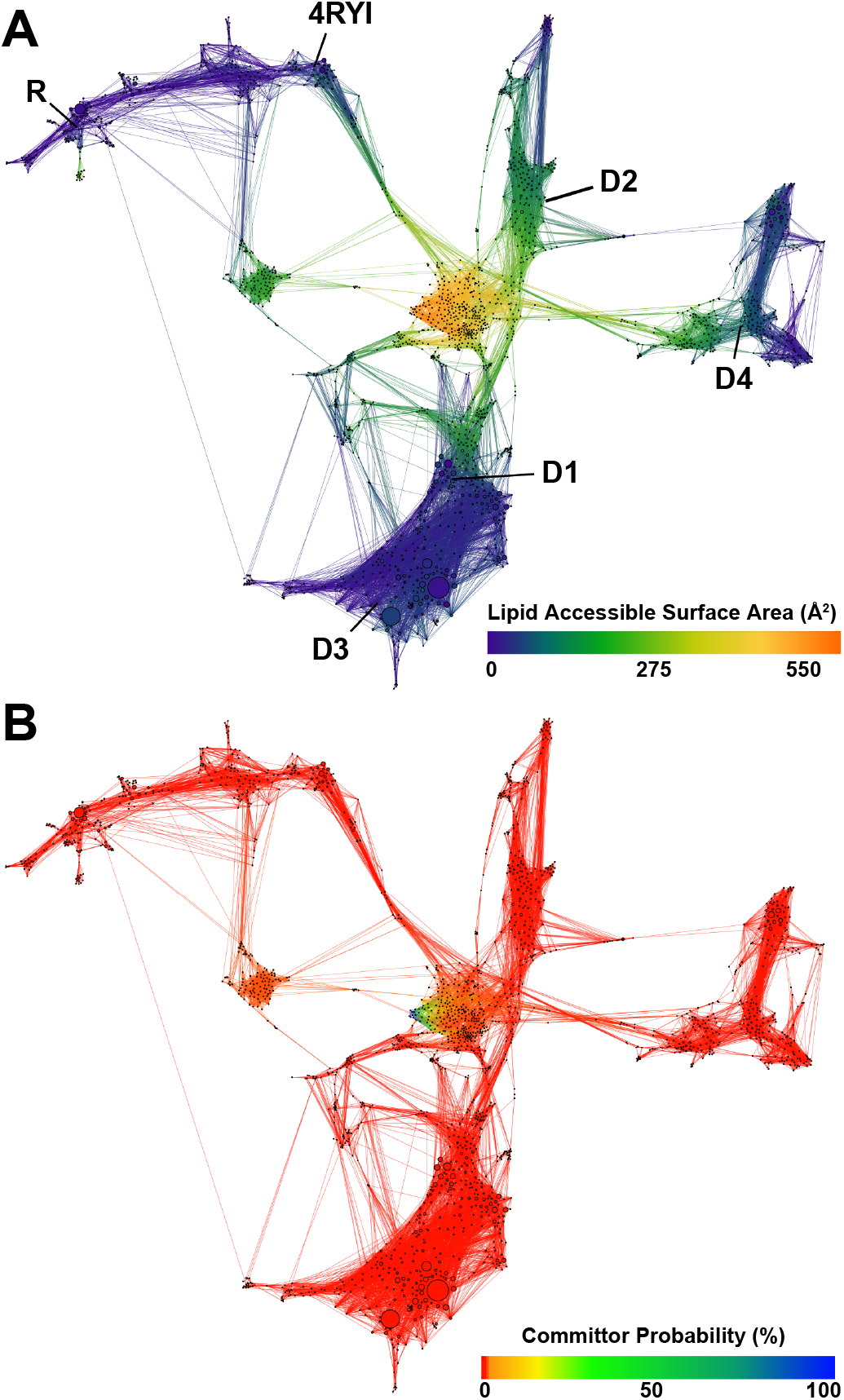
Combined conformational space network of all REVO simulations from each starting pose with the addition of frames from straightforward MD simulations, colored by (A) LASA and (B) committor probability. Starting poses are marked in bold in A.

With the addition of the straightforward simulations, we recalculated the MFPTs from each starting pose (Fig. 4C). The MFPTs of all poses are now nearly identical at low lag times. This indicates that with the additional information gained from the straightforward simulations, the poses can all interconvert before unbinding. By again tracking the probability flow of each pose during the Markov chain simulation, we see that pose D1 and D3 have the highest probability before unbinding regardless of the starting pose. (Fig. S13). Additionally we observe the MFPT curve in Fig. 4C is smoother at low lag times than the CSN with only the REVO simulations. This suggests the transition matrix is less sensitive to the addition and removal of nodes and pathways at lag times below 2.25 ns. Between lag times of 0.8 and 1.5 ns, the MFPT is at its most consistent and the predicted MFPT results are on the same order of magnitude as those determined experimentally (2). At higher lag times the MFPT calculation fluctuates wildly similar to the results of the MFPT on the REVO-only CSN network.

Fig 5B, shows the committor probability of each state in the final network (Fig 5). The vast majority of the states have a near zero committor to the unbound state. Only once PK is dissociated into the membrane does the committor probability begin to significantly increase. We find that the transition state – where the committor equals 0.5 – occurs when PK has dissociated into the membrane and is at a minimum distance of least 4 Å away from TSPO. For these states we find that the non-bonded interaction energy between PK and TSPO is negligible, below 1 kcal/mol (Fig. S10). This suggests that the membrane presents a physical barrier that acts to trap PK near TSPO when it initially dissociates into the membrane.

## DISCUSSION AND CONCLUSION

The results of our simulation show that from all six initial PK poses, using the *R. sphaeroides* TSPO structure, the ligand dissociates into the membrane through the transmembrane helices. We found a pathway between TM1 and TM2 and a lower probability pathway between TM2 and TM5. These pathways identify residues with which PK has high interaction energy. Among them are aromatic residues: Phe46 and Trp50 which form *π*-*π* interactions with the ligand. The interactions with the Trp50 rings are also found in different bound states. We note that the Trp50 residue happens to be highly conserved across organisms of several species and kingdoms. These stabilizing interactions could lower the barrier to entry for other TSPO ligands such as protoporphyrin-IX and heme, which are also largely aromatic.

Previous results (18) using a different starting pose and TSPO structure showed PK dissociating into the cytosol through the LP1 loop region. The TSPO structure used in the previous study was built from a homology model based on the mouse NMR TSPO structure and used the rat sequence, whereas our structure was determined from X-Ray crystallography from *R. sphaeroides* TSPO. As mentioned in the Introduction, this NMR structure was destabilized by the detergent used in the purification (21, 22), which likely affected the homology model structure as well. This, in addition to the differences in sequence, results in several key structural differences between the mouse (PDB 2MGY (6)) and *R. sphaeroides* (PDB 4UC1 (4)) structures. TM1 in the mouse structure is significantly longer and the top portion of the helix is at a drastically different angle than the helix in the structure we used in our simulations. While the LP1 region is present in both structures, the *R. sphaeroides* sequence has a small *α*-helix which in the mouse structure is incorporated into TM1. Finally, the LP1 region in *R. sphaeroides* has several stabilizing interactions (4) between non-bonded residues such as between Trp30-Met97, Asp32-Arg43 and Trp39-Gly141 that are not present in the mouse structure. This stabilization limits the freedom of motion of the LP1 loop, sterically hindering PK from leaving via the LP1 pathway. Additionally, previous results were obtained using a 2:1 POPC:cholesterol lipid bilayer, while our results used an approximately 2.9:1.6:1 mixture of POPC:POPE:POPI lipids. Cholesterol is known to bind to TSPO, although known binding sites are not close to the TM1-TM2 pathway found here. Differences in lipid composition could also affect membrane fluidity, which could impact the relative probabilities of the LP1 and TM1-TM2 pathways. A third difference between the two studies is the type of enhanced sampling method used to generate the unbinding pathways. Bruno et al (18) used the random acceleration molecular dynamics method (19) to generate their initial paths, which were subsequently examined using steered molecular dynamics (20). It will be an important goal of future work to parse the relative impact of these four differences (protein sequence, protein structure, membrane composition and sampling methodology) in determining ligand dissociation pathways.

There is interest in designing new TSPO ligands with longer residence times (2, 17). The ligand binding transition state is the rate-limiting step of ligand binding and release, which can also be identified in simulations by a committor probability of 0.5 between the bound and unbound basins. Here we find that the ligand binding transition state occurs when the ligand is no longer in direct contact with TSPO, but has approximately 4 Å of clearance between the closest ligand and protein atoms, and has minimal interaction energy between the protein and ligand. In addition to details of the bound state, this implies that TSPO ligand residence time is primarily affected by properties related to membrane permittivity and diffusivity, such as hydrophobicity. These results lead to the hypothesis that the membrane composition could have a direct impact on ligand binding kinetics of PK.

This work also raises questions about membrane insertion and removal along ligand binding paths. Additional REVO simulations with only PK and the lipid membrane could reveal the membrane diffusion coefficient of PK as well as rate constants for insertion and removal to form holistic models of membrane-mediated binding from solvent to binding site. A larger question is how the presence of other proteins known to interact with TSPO, such as VDAC (voltage dependent ion channel) (9) and cytochrome P450s (40) affect the unbinding/binding and insertion/removal pathways. Cholesterol could also affect the binding pathways of PK, either by binding to TSPO and affecting a conformational change, or through membrane fluidity, which could affect the (un)binding rate of PK as it interacts with the membrane (41).

One thing to note is the experimental MFPT reported by Costa (2, 17) was determined using human TSPO while our simulations used the structure from *R. sphaeroides* containing a A135T mutation. While the mutation was designed to mimic the human TSPO structure (4), the human and *R. sphaeroides* have low sequence homology (30%) which could result in different transition paths, transition states and unbinding rates. A more precise comparison would be to the MFPT of PK from the *R. sphaeroides* A135T mutant, which we predict would be similar or slightly longer than the MFPT for human TSPO. Furthermore, these results emphasize that we should take care to ensure consistency of the “unbound” state from simulation and experiment. In radioligand displacement assays, any ligand pose that is not sterically blocking entry of the radiolabelled competitor ligand would be considered “unbound” (42). However, in surface plasmon resonance, a ligand would still be considered bound until it dissociated from the detergent that is bound to the chip along with TSPO. Our simulations show how differences in the definition of the unbound state can lead to significant differences in residence time, and could help rationalize differences between experimental residence times obtained with different methods.

## AUTHOR CONTRIBUTIONS

T.D., S.F.M. and A.D. designed the project. T.D. and A.U. prepared and conducted the simulations. T.D. analyzed the data. T.D. and A.D. wrote the manuscript, which was edited by all the authors.

## ACKNOWLEDGEMENT

This work was supported by R01GM130794 from the National Institutes of Health. The authors thank Dr. Jens Meiler and Georg Kuenze for providing the pose R structure from Ref. (32).

## SUPPLEMENTARY MATERIAL

An online supplement to this article can be found by visiting BJ Online at http://www.biophysj.org.

