## Supplemental Information for "Membrane-mediated ligand unbinding of the PK-11195 ligand from TSPO"

Manuscript submitted to **Biophysical** *Journal*
Supplementary Material

### Membrane-mediated ligand unbinding of the PK-11195 ligand from TSPO

Tom Dixon[1,2], Arzu Uyar[2], Shelagh Ferguson-Miller[2], and Alex Dickson[1,2,*]

[1]Department of Computational Mathematics, Science and Engineering, Michigan State University, East Lansing, USA
[2]Department of Biochemistry & Molecular Biology, Michigan State University, East Lansing, USA

#### S1 SUPPLEMENTAL METHODS

##### S1.1 Markov Chain Simulation Probability Tracking

In addition to tracking the probability in the unbound state during the Markov chain simulations, we also tracked the probability in each starting basin. When a cluster was visited by multiple starting poses during the MD simulations, we assigned it to the basin the highest weighted walker originated from. Due to the high connectivity between the D1 and D3 starting poses, we combined those two basins into the D1-D3 basin. We performed these calculations both for the networks with (Fig. S13) and without (Fig. S12) the straightforward MD simulations from each starting pose.

#### S2 SUPPORTING FIGURES

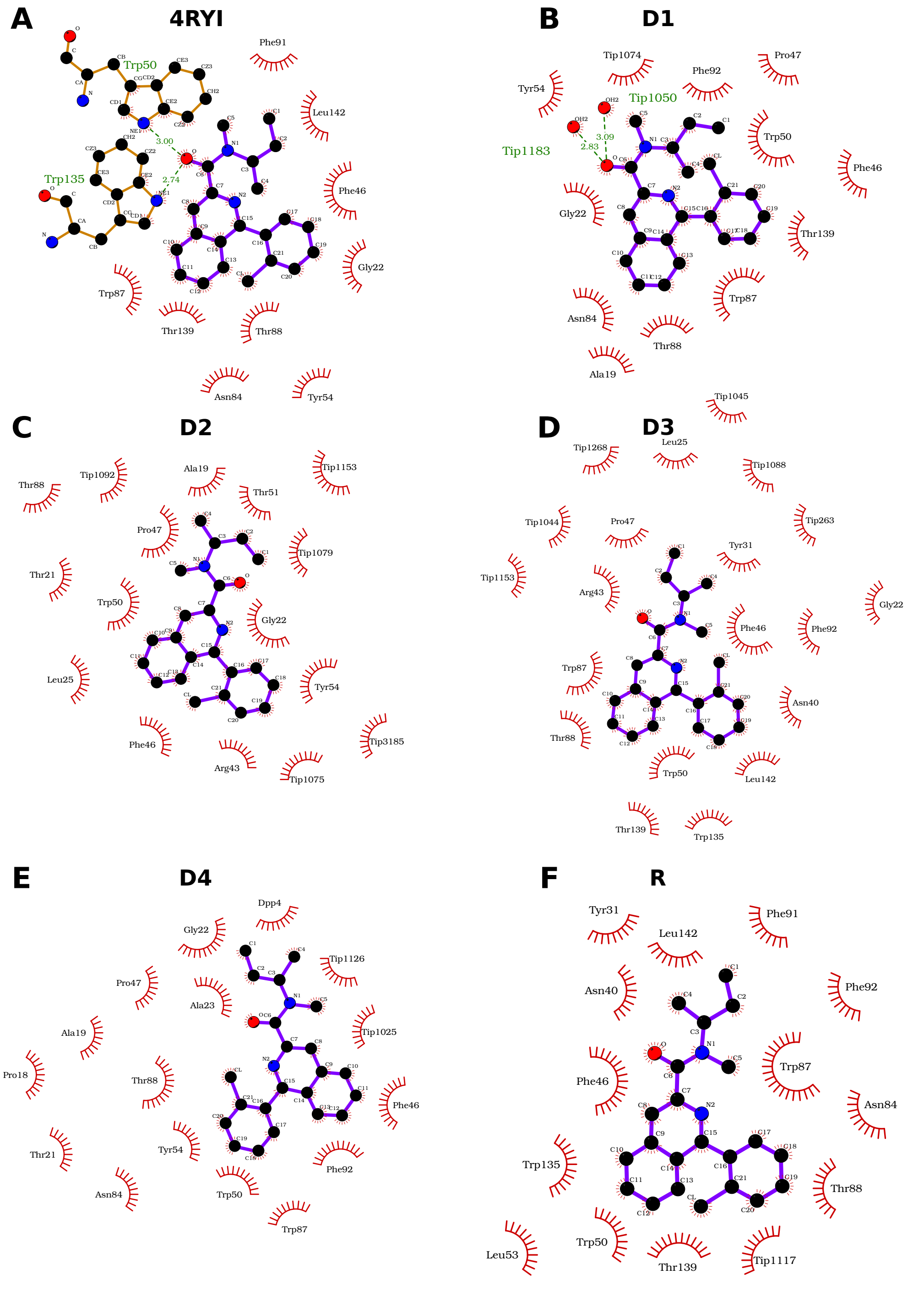


Figure S1: Protein-ligand interaction plots for the six starting conformations. The red suns indicate that the residue has a hydrophobic contact with PK. The green dashed lines show hydrogen bonds.

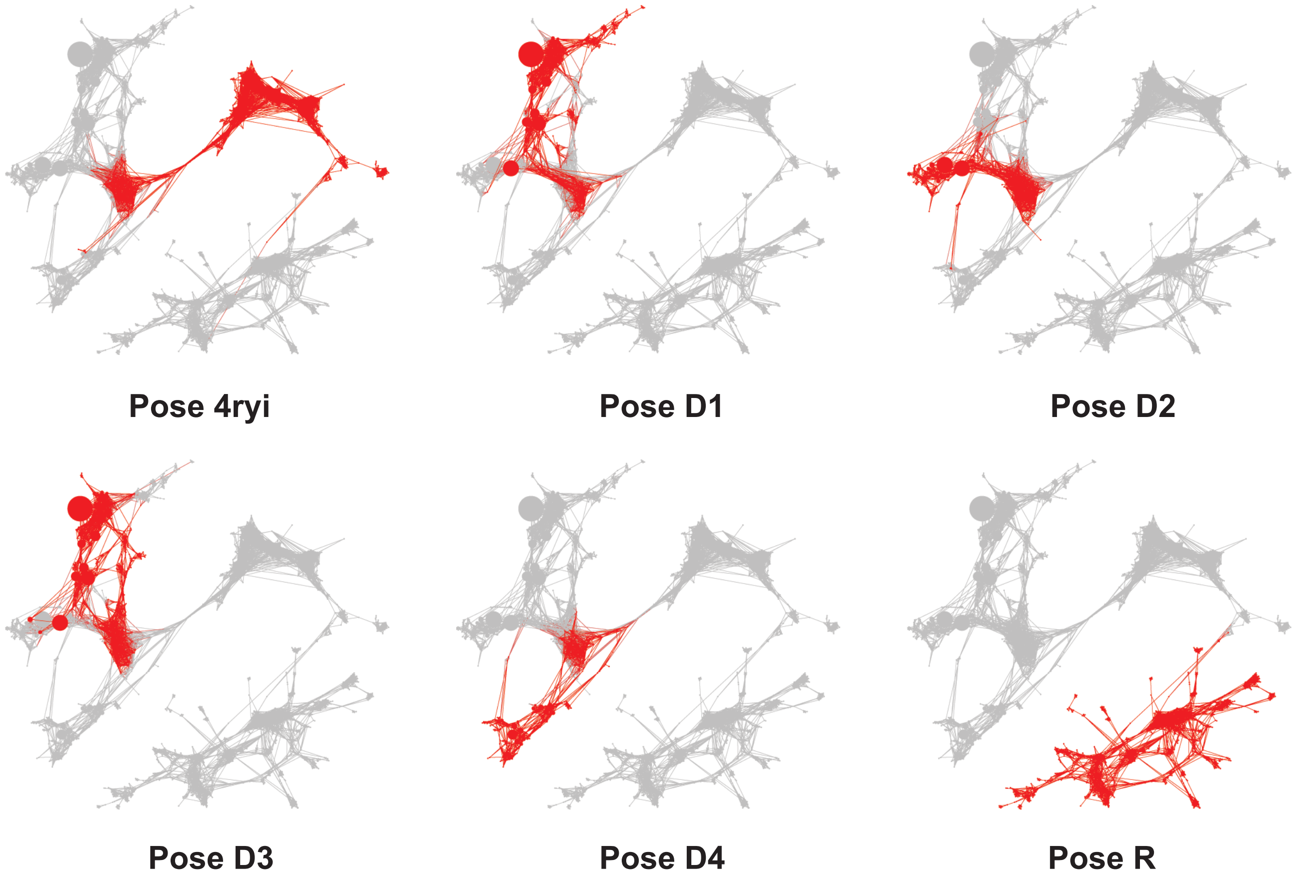


Figure S2: CSN networks indicating the clusters that were observed from each initial pose. Red nodes indicate the simulations observed a TSPO-PK conformation that was clustered into that node.

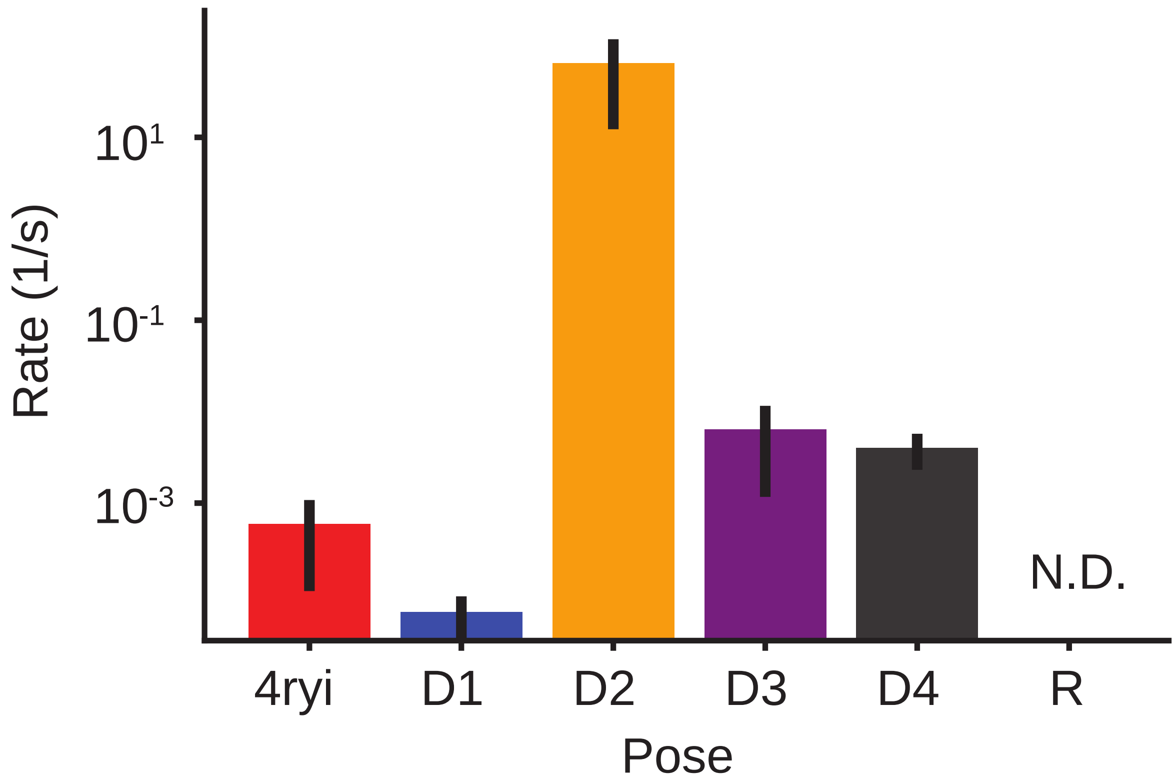


Figure S3: Predicted rates for each of the starting poses for a 5 Å minimum distance between PK and TSPO. The error bars are the standard error for the rates of each pose. PK never met that boundary condition to calculate a rate for pose R, leaving that rate undetermined.

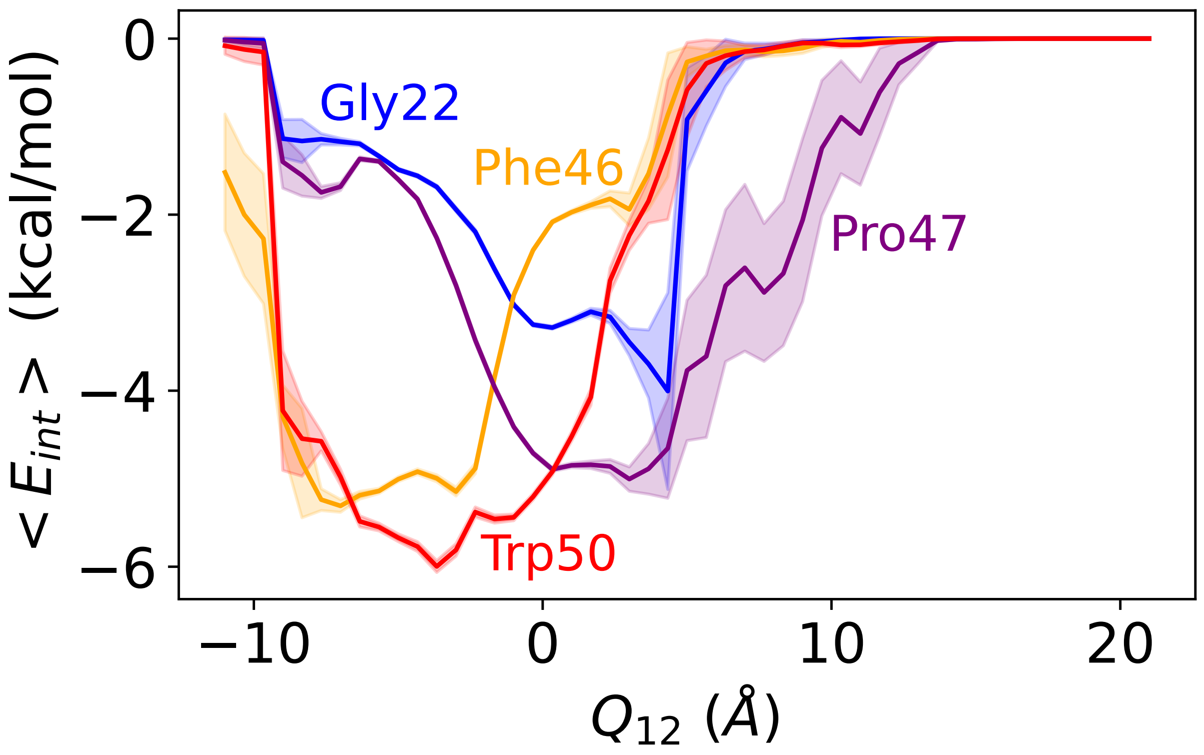


Figure S4: Expectation value for $E_{int}$ as a function of $Q_{12}$. The lines are colored by residue. Only residues who have a minimum interaction energy below −3.5 kcal/mol are shown. The standard error is shown in the lighter shaded regions.

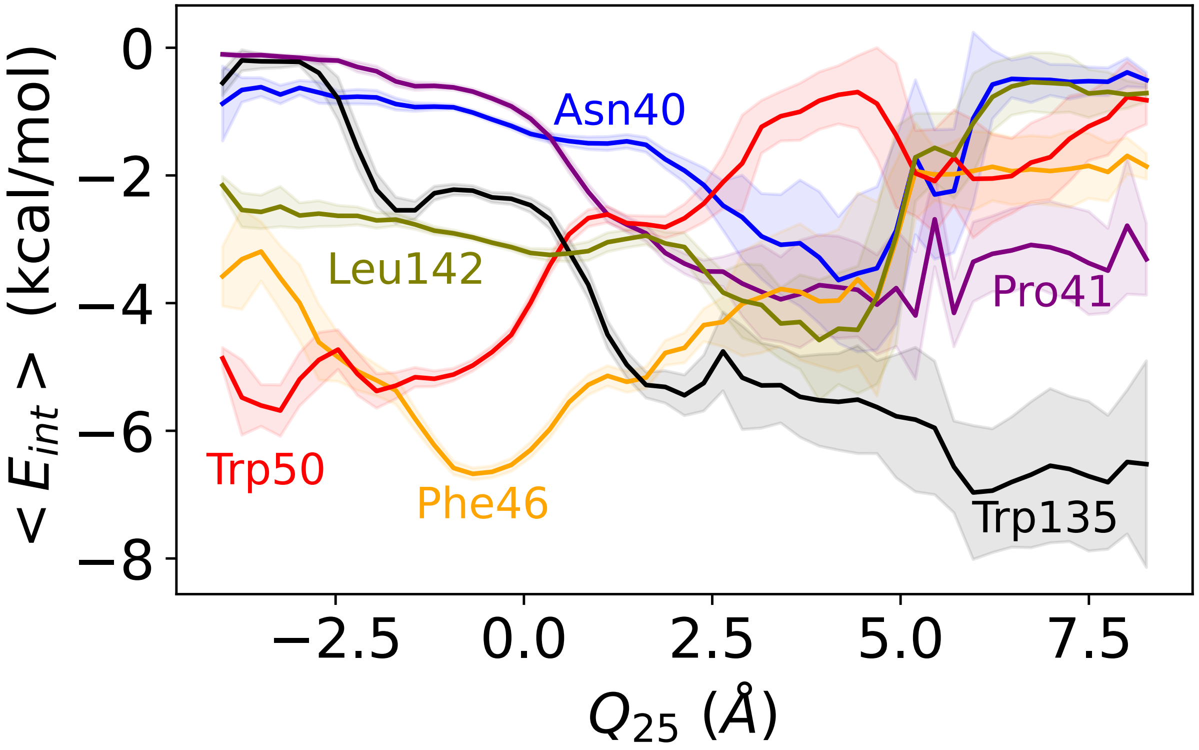


Figure S5: Expectation value for $E_{int}$ as a function of $Q_{25}$. The lines are colored by residue. Only residues who have a minimum interaction energy below −3.5 kcal/mol are shown. The standard error is shown in the lighter shaded regions.

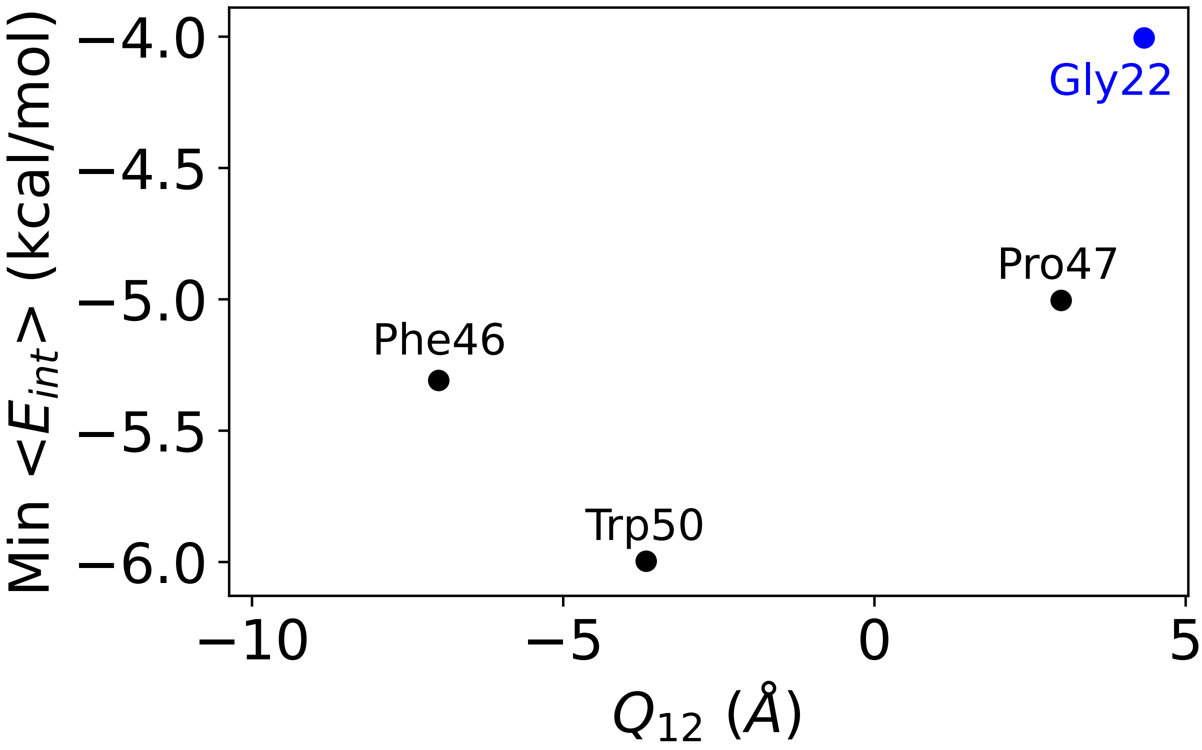


Figure S6: The residues with the strongest non-bonded interactions with PK on the $Q_{12}$ pathway. This summarizes the curves in Fig. S4, plotting the minimum $E_{int}$ against the $Q_{12}$ value for which this minimum value is observed. The colors indicate the region of TSPO, blue for residues on TM-1 and black for residues on TM-2. Only residues with a non-bonded energy below -3.5 kcal/mol are shown.

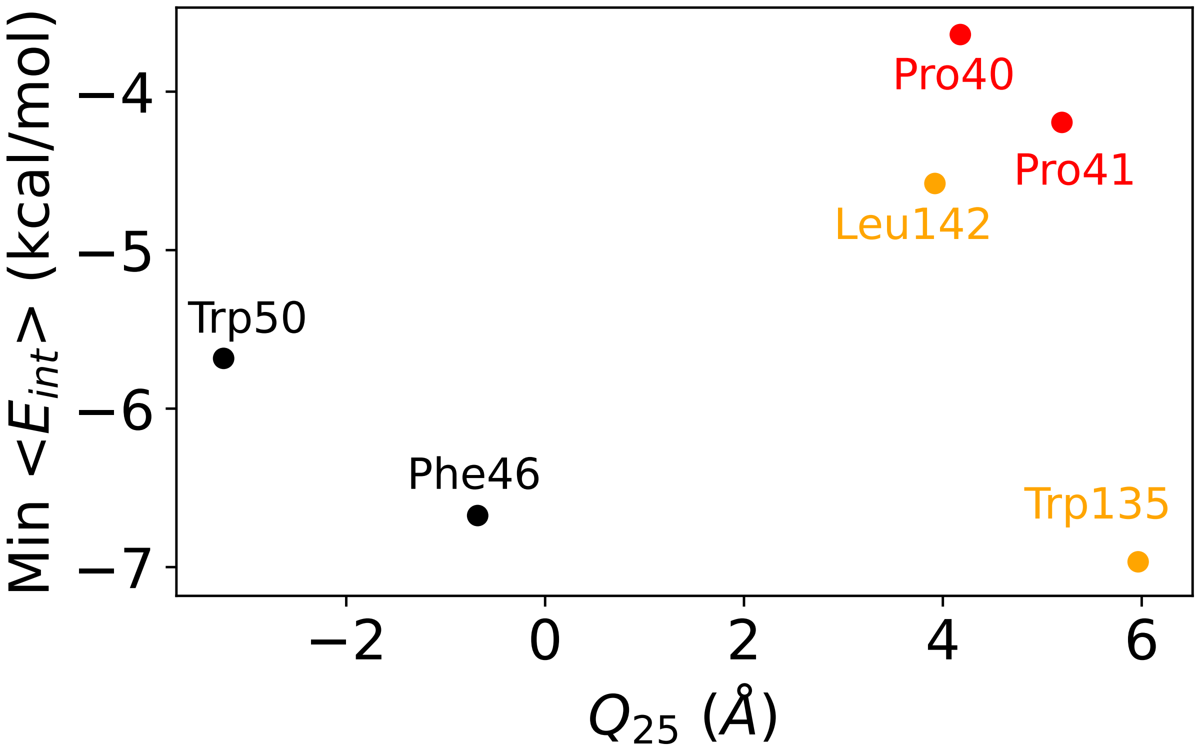


Figure S7: The residues with the strongest non-bonded interactions with PK on the $Q_{12}$ pathway. This summarizes the curves in Fig. S5, plotting the minimum $E_{int}$ against the $Q_{25}$ value for which this minimum value is observed. The colors indicate the region of TSPO, red indicates residues on the LP1 loop, black for residues on TM-2 and orange for residues on TM-5. Only residues with a non-bonded energy below −3.5 kcal/mol are shown.

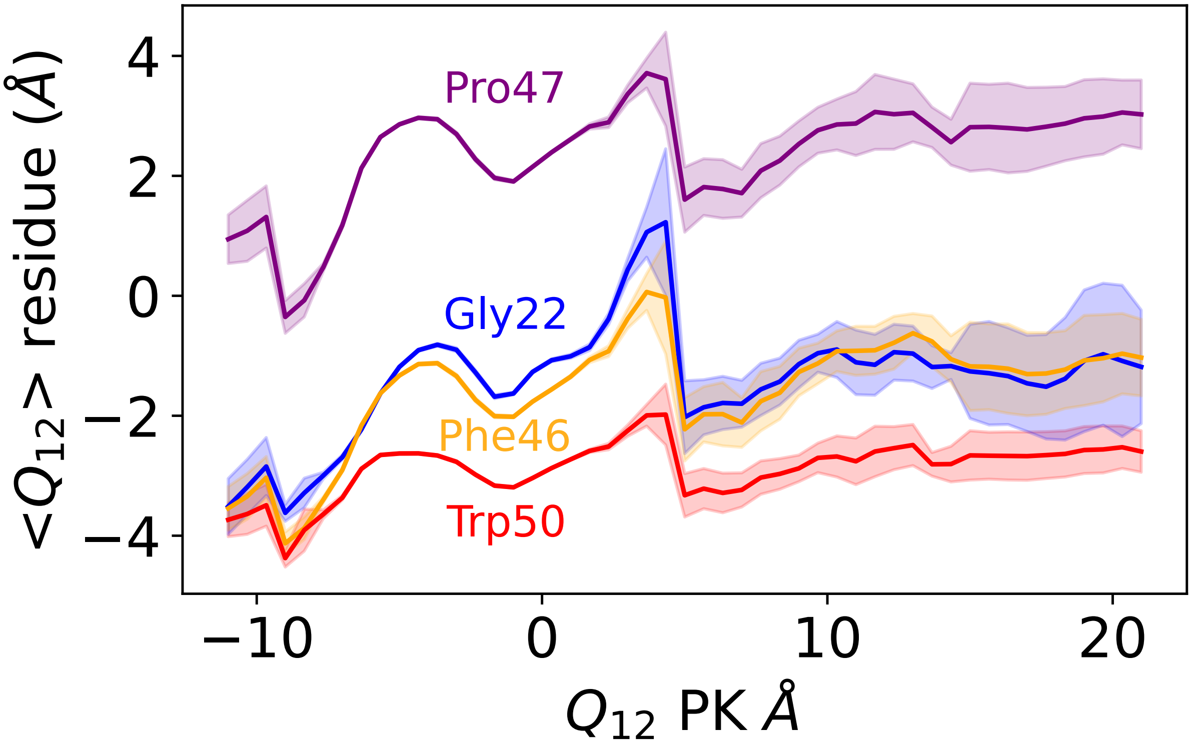


Figure S8: Residues moving along with the ligand during dissociation. Expectation values of $Q_{12}$ for individual residues are shown as a function of the $Q_{12}$ of PK.

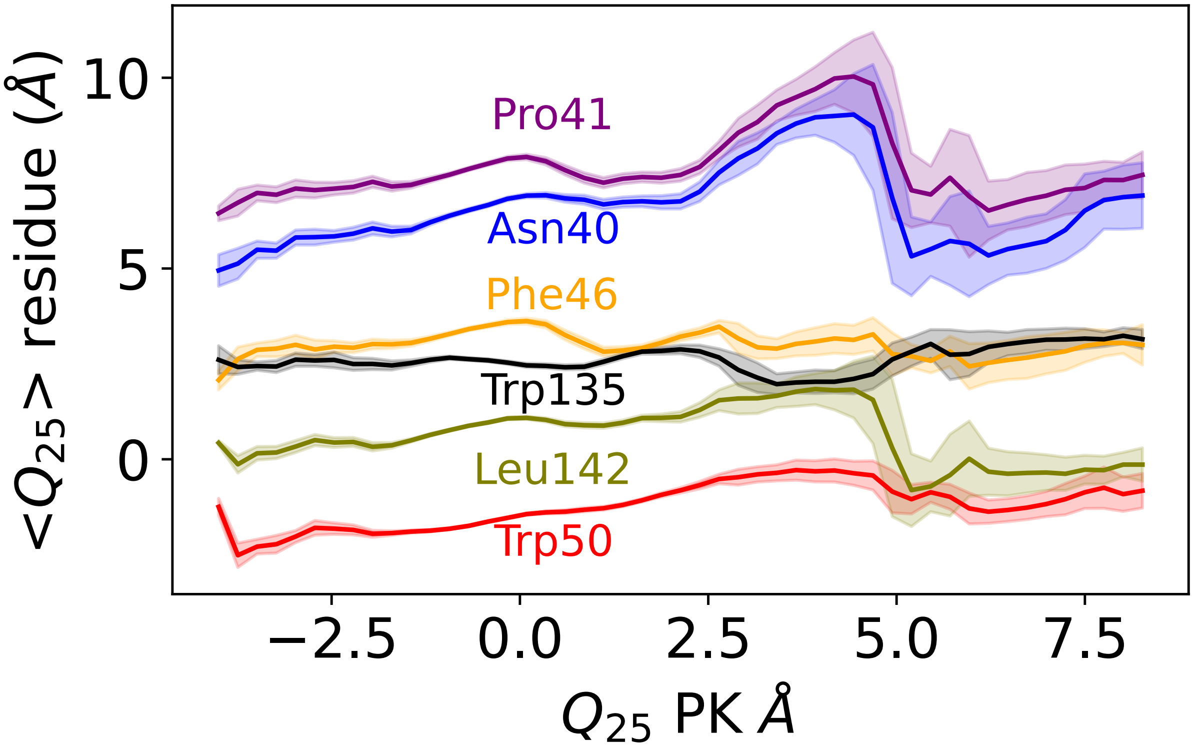


Figure S9: Residues moving along with the ligand during dissociation. Expectation values of $Q_{25}$ for individual residues are shown as a function of the $Q_{25}$ of PK.

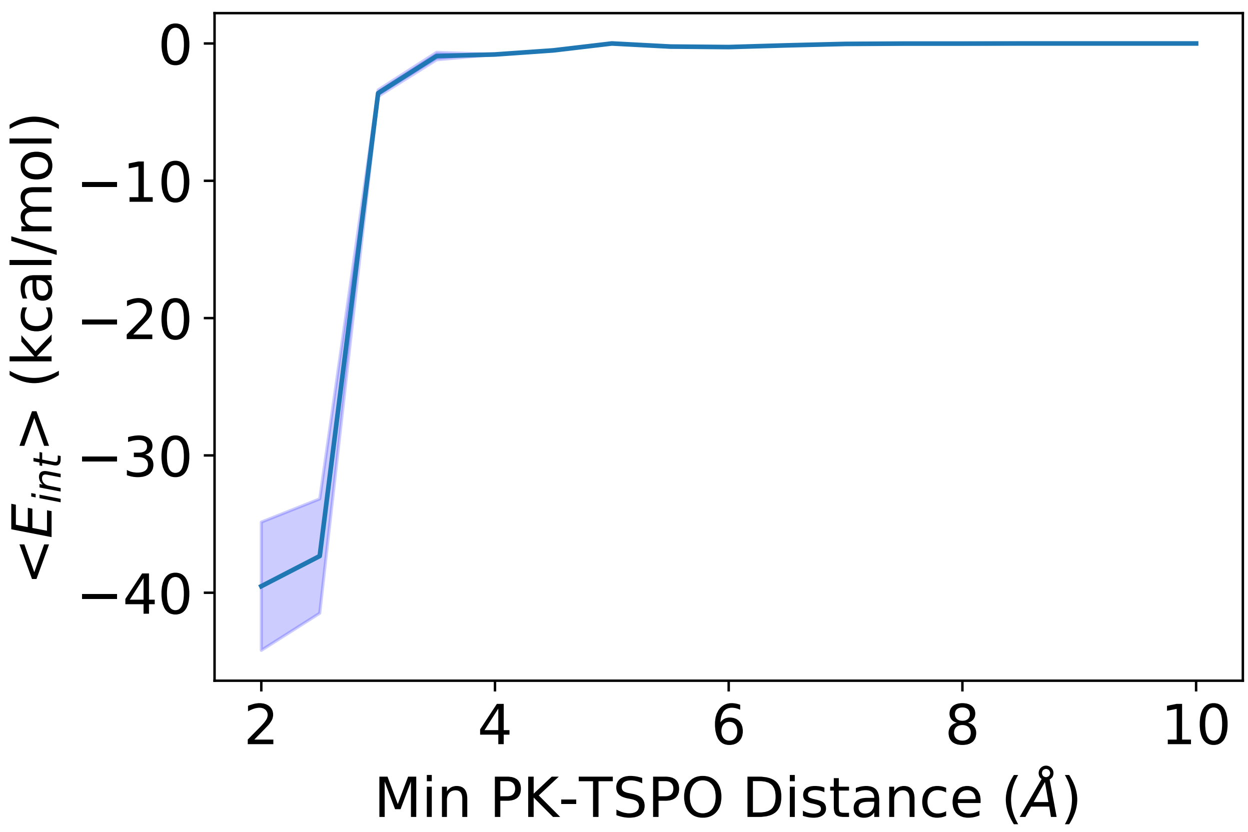


Figure S10: The energy of non-bonded interactions between PK and TSPO as a function of minimum distance between PK and TSPO.

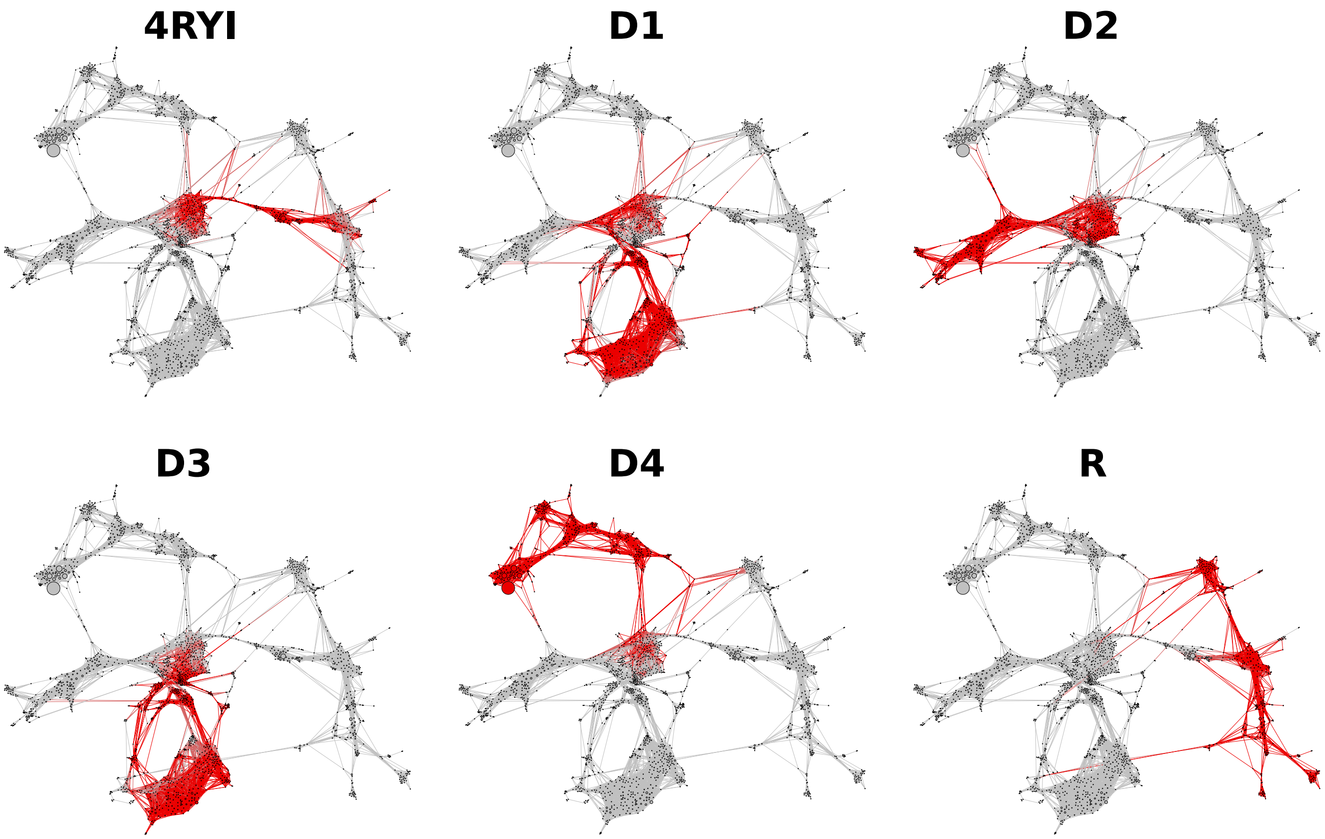


Figure S11: CSN networks, including both the straight forward and REVO trajectories, indicating the clusters that were observed from each initial pose. Red nodes indicate the simulations observed a TSPO-PK conformation that was clustered into that node.

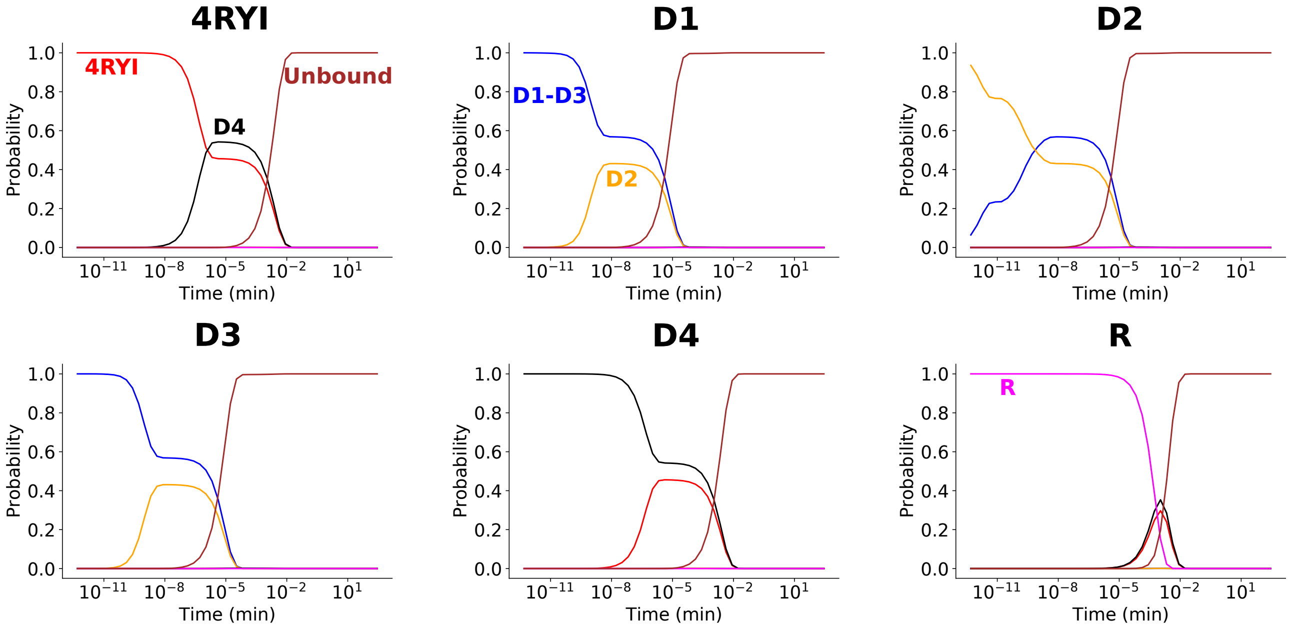


Figure S12: The probability of being in any given basin over the course of the Markov chain simulations for the REVO only network. Each graph represents starting the simulation with all the probability at a specific starting pose.

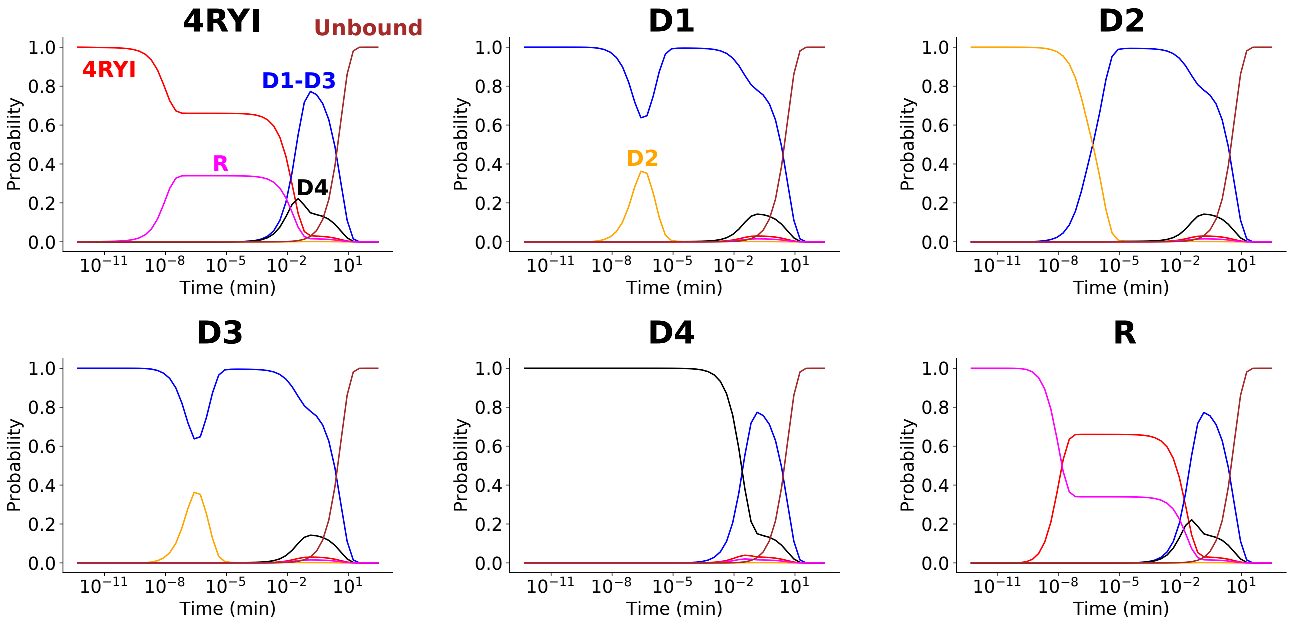


Figure S13: The probability of being in any given basin over the course of the Markov chain simulations for the network containing REVO and straightforward simulation frames. Each graph represents starting the simulation with all the probability at a specific starting pose.
